# Anticipated action goals structure spatial organisation in visual working memory following self-movement

**DOI:** 10.1101/2025.05.01.651644

**Authors:** Babak Chawoush, Dejan Draschkow, Freek van Ede

## Abstract

Working memory enables the retention of relevant visual information in service of anticipated behaviour. Consequently, the goal for which information is retained may crucially sculpt the way we retain and organise information in visual working memory. Here, we investigated how anticipated action goals structure the spatial organisation used for visual working memory following self movement. Participants encoded two tilted objects (left and right) in working memory, then turned around 180 degrees before being cued to either reproduce the tilt of the cued object from memory (report session) or manually reach back to it (reach session). The 180-degree self-movement uniquely enabled us to oppose and disentangle two potential spatial frames for immersive working memory in moving participants: the native spatial frame of how the objects were registered during encoding and an updated spatial frame keeping track where the objects are in the external world behind the participant after self-movement. Behavioural memory reports and spatial biases in gaze converged on the use of distinct spatial frames in the two tasks. This reveals how a foundational aspect of working memory in behaving humans – the spatial frame used to organise memories following self movement – is critically sculpted by anticipated action goals.

## Introduction

Working memory enables us to carry forward recently encountered visual information in service of anticipated upcoming behaviour ^1–4^. For example, while momentarily turning away from your dance partner, relevant visual information may become out of sight, but can be held ‘in mind’ (in working memory) to ensure a smooth interaction with your dance partner upon returning. This example underscores two foundational aspects of working memory. First, it underscores its future-focused nature: we keep in working memory those aspects of our environment that we anticipate to become relevant for ongoing and upcoming behaviour. Second, it underscores how we often engage working memory by keeping in mind aspects of our environment that our own behaviour renders out of sight, as we ourselves move through the world.

Anticipated behavioural demands can shape not only ‘what’ visual information we decide to bring into working memory, but also ‘how’ we retain this visual information in working memory, such as the format in which information is maintained and the brain areas and networks that are recruited (e.g., ^5–13^). This is the case, even when the visual information available at encoding is identical. For example, when the same visual objects are anticipated to become relevant for a task requiring precise identity or more generic category information, distinct brain areas are recruited to maintain the information in working memory in a goal-dependent format ^12^. Likewise, when we expect to be tested on a certain visual feature of an object in memory (e.g. its colour or orientation), we retain the relevant visual feature with greater priority ^14,15^, and recruit feature-specific brain circuitry ^11^.

Besides shaping the retention of specific visual features and the invocation of specific neural networks, the anticipated task may also fundamentally shape the way in which such visual objects are *organised* in working memory. One foundational organising principle in visual working memory is the use of space (memorised object locations) as a scaffold. Indeed, ample studies have shown how space serves to individuate, rehearse, and select visual objects in working memory, even when spatial location is never asked about (e.g., ^16–27^).

The organising role of space for working memory becomes particularly interesting when considering working memory in actively behaving participants that move through the three-dimensional environment. Unlike conventional laboratory tasks of visual working memory in which participants remain seated and memorise static objects presented on 2D displays, when moving around in the three-dimensional world, we may be prompted to update spatial representations and remember information with regard to multiple spatial frames ^28–37^. In a virtual-reality (VR) study with moving participants, we recently brought the study of spatial reference frames to the domain of visual working memory, revealing how the spatial organisation of visual working memory may rely on multiple spatial frames ^38^ (see also: ^39–42^). This was the case even though memorised object locations were never explicitly asked about ^38^, underlining the vital role of space in memory organisation. Thus, when moving around the world, we may rely on multiple spatial frames to organise the contents of visual working memory. Building on this premise, we here hypothesise that the predominant spatial frame used for organizing visual working memory following self-movement will be fundamentally sculpted by the anticipated memory task – even when retaining identical visual information.

To address this hypothesis, we developed a VR task in which participants encoded two oriented bars (encoded to the left and right of fixation), then turned around (rotated 180 degrees) before being cued to either reproduce the memorised orientation of the cued object (report task) or manually reach back to it (reach task). The 180-degree participant rotation uniquely enabled us to oppose and disentangle two potential spatial frames used for working memory organisation: the native spatial frame of how the objects were registered during encoding and an updated spatial frame keeping track where the objects are in the external world behind the participant after self-movement. We were able to do so by investigating the distributions of orientation reports (that, after rotation, are mirror images in the two frames) as well as through horizontal left/right biases in gaze after the memory cue (cf. Chawoush et al., 2023; de Vries et al., 2023; Liu et al., 2022; van Ede et al., 2019; Wang & van Ede, 2024) that are predicted to go in opposite directions in the two frames. Doing so, we provide two independent and corroborating sources of evidence that the spatial organisation used for visual working memory following self movement is fundamentally sculpted by anticipated memory-guided behaviour, i.e., by the purpose for which we memorise.

## Results

Participants performed an immersive visual working-memory task in VR (**Fig. 1**). In different sessions, participants either reproduced the precise orientation of the cued visual object (report task), or reached back to grab it (reach task); after having turned their backs to the object following a 180-degree rotation. Importantly, following 180-degree rotation, memory objects could be represented in two distinct spatial frames: in a native encoding-centred frame of how the objects were registered before rotation (an eye-centred picture-like snapshot representation), or in an updated body-centred frame keeping track where the visual objects are in the external world behind the participant following self-movement. The critical innovation here is that following 180-degree rotation, these two frames are associated with (1) mirror-symmetric memorised object tilts (“\” vs. “/”) and (2) opposite memorised object directions (left vs. right). Accordingly, we could interrogate the spatial frame used for immersive visual working memory by inspecting: (1) the reported object tilt and (2) the left/right direction of spatial biases in gaze during mnemonic selection. These two sources of evidence yielded converging evidence in support of goal-dependent use of distinct spatial organisations for working memory following self-movement.

**Figure 1.**
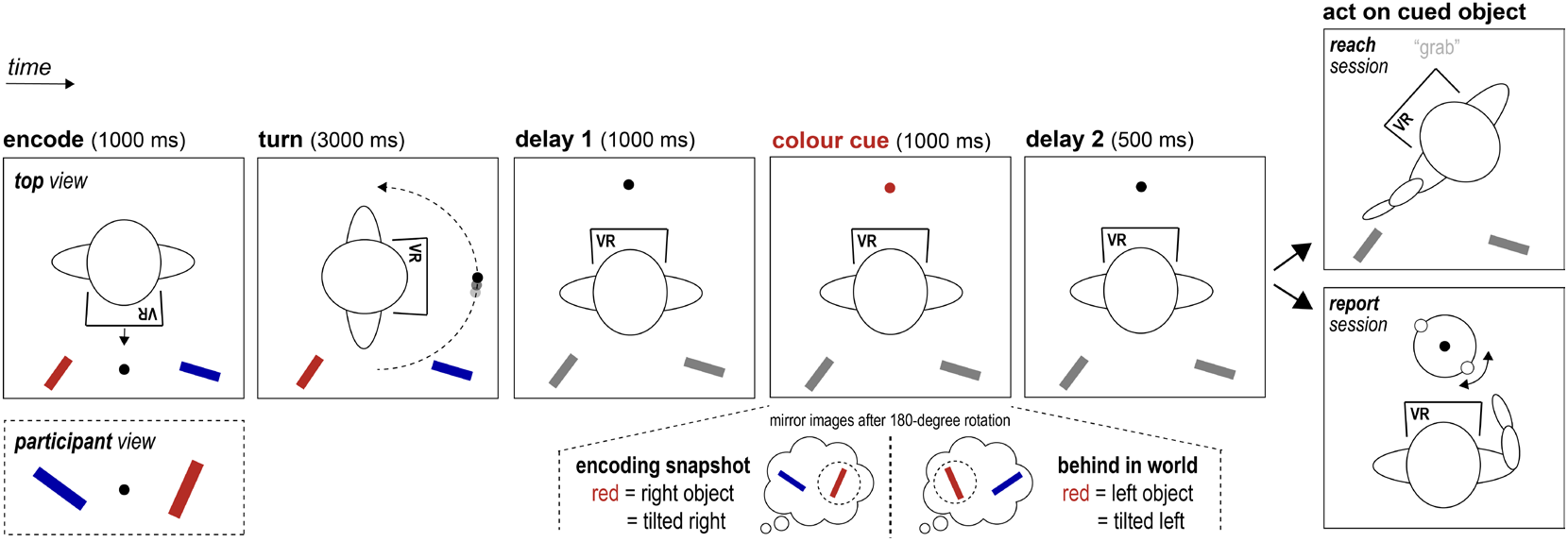
Schematic of experimental task and logic. Participants performed an immersive visual working-memory task in Virtual Reality (VR). Participants encoded two titled visual cylinders presented to left and right at encoding, then turned around 180 degrees. After rotation, a colour change of the central fixation marker cued which object – now behind the participant and thus in working memory – would become tested moments later. In different sessions, participants either reproduced the precise orientation of the cued visual object (report task) or reached back to grab it (reach task). The inset at the bottom shows how following a 180-degree rotation, memory objects could be represented in two distinct spatial frames that were associated with mirror-image orientations and opposite left/right locations (see text for further elaboration).

### Orientation reports reveal the use of a snapshot-like representation when memorising for reporting

To successfully perform the reach task, participants were naturally required to represent where the memory objects are in the world behind them. Indeed, if participants would rely exclusively on how the objects were registered prior to turning around, then they would consistently turn back in the wrong direction. This is because the object registered to the left before rotation (in retinal coordinates) would, after rotation, require reaching back to the object that would now be to the right behind the participant in the external world. Consistent with this assertion, participants rotated back in the correct direction in 95.553±0.006% [M±SEM] of all reach trials. This verifies the use of spatial representations for visual working memory that were updated from their native encoding (retinotopic) frame, by incorporating where the objects currently reside in the world behind the participant (relative to one’s current facing direction, or body-centred).

Critically, in contrast to the reach task, in the complementary report task, there was no a-priori priority for the use of one frame over another. Participants were simply asked to reproduce the precise orientation of the cued memory object (while remaining in their current facing direction after rotation). This task could potentially be solved by memorising the objects in at least two distinct ways. Imagine encoding an object with a leftward tilted orientation (“\”). After turning around 180 degrees, you may remember this object precisely as you have seen it (“\”; i.e. as if carrying a picture of it with you), or update your representation to match the object’s orientation in the world behind you, which is now tilted in the opposite way (“/”) relative to your current facing direction. In other words, after a 180-degree rotation, objects could be remembered precisely as they were registered during encoding (eye-centred), or as their exact mirror image (body-centred, updated to match their orientation in the world behind the participant). Because we never explicitly informed participants that there were these two potential frames to report the orientation (nor that one frame was more “correct” than the other), we could uniquely use the orientation reports in this task to interrogate which spatial frame participants naturally or spontaneously relied on.

Orientation reproductions in report blocks consistently concentrated around the orientation of the cued memory object as it was perceived during encoding (**Fig. 2a**) rather than being updated to represent the object orientation as imagined in the world behind the participant. This was quantified by comparing the average absolute deviation from the target orientation relative to both potential spatial frames (**Fig. 2b**). Consistent with the rich visualisation from the density plots (**Fig. 2a**), this revealed a clear preference for reports to follow the picture-like snapshot representation, with errors being lower when calculated relative to the orientation as perceived prior to rotation (**Fig. 2b**; t(49) = 6.724, p < 0.001, d = 0.951).

**Figure 2.**
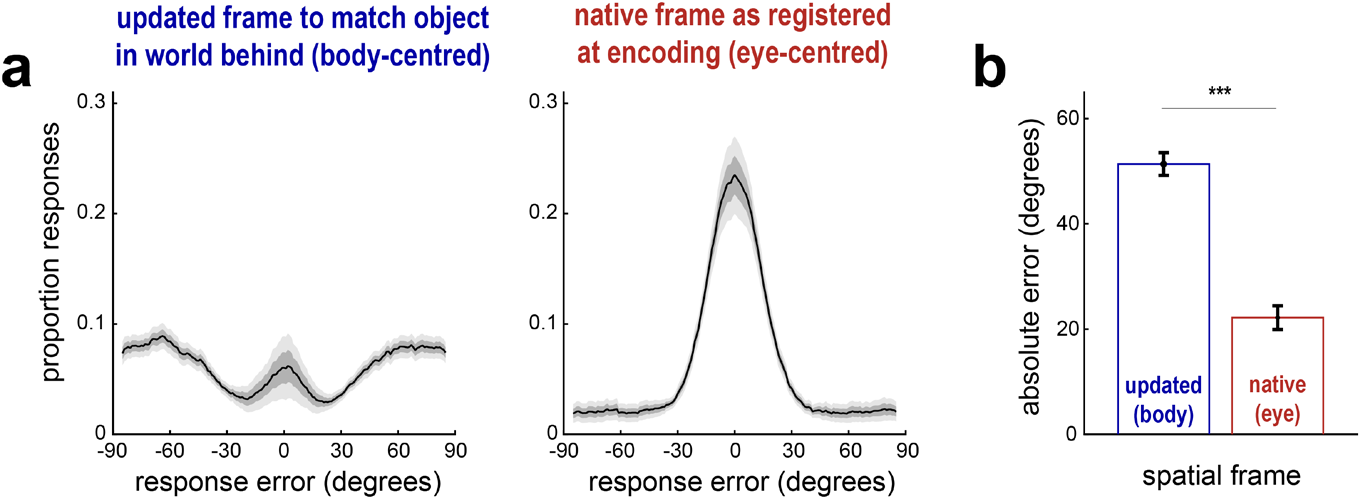
Orientation reports reveal the use of a snapshot-like (eye-centred) representation when memorising for reporting. **a)** Response densities for orientation-reproduction responses in the report task. The left panel shows reproduction errors relative to the cued memory object as defined in the updated spatial frame that matches how the object is tilted behind the participant (body-centred). The right panel shows errors defined relative to the orientation of the cued memory object as registered during encoding, prior to rotation (eye-centred). **b)** Trial-average absolute reproduction errors as defined in both spatial frames. *** p < 0.001.

These data alone already suggest the use of distinct spatial frames used for working memory organisation in the two tasks. In the next section, we provide additional evidence from spatial biases in gaze during mnemonic selection that reinforced the use of distinct spatial frames used for the two tasks – despite relying on the same memorised visual information.

### Spatial biases in gaze during mnemonic selection corroborate distinct spatial frames when memorising for reaching versus reporting

We next turned to a complementary signal that we measured at an earlier stage in the trial and that we could uniquely asses in both tasks: directional biases in gaze signalling mnemonic selection within the spatial-layout of working memory (building on: ^17,25,43–45^). The logic was as follows: if the spatial organisation in memory engages distinct frames when retaining both objects after rotating, then gaze signatures of mnemonic selection should go in *opposite* directions. To appreciate this, consider again the object that is to the left of the participant at encoding. In the native eye-centred frame – representing a picture-like snapshot of the objects as registered during encoding – the object remains represented as the left object, even after rotation. In contrast, in the updated body-centred frame – keeping track of where the object is in the world behind the participant relative to the current facing direction – the left object at encoding becomes represented as the object to the right behind the participant.

To expose whether participants considered objects to the left or right during mnemonic selection – without explicitly asking participants – we contrasted gaze-position densities between trials in which the central colour cue (presented after rotation), cued the object that was left (blue object in **Fig. 3**) or right (red object in **Fig. 3**) during encoding. This is depicted in **Figure 3** where the difference in gaze density is colour-coded, and where the x-axis shows the horizontal (left-right) axis of gaze, while the y-axis shows time after the retrocue. The difference in gaze density is coded in such a way that red shows where participants looked more when selecting the red object and blue shows where participants looked more when selecting the blue object (with “red” and “blue” being defined in reference to the example configuration in **Fig. 3**).

**Figure 3.**
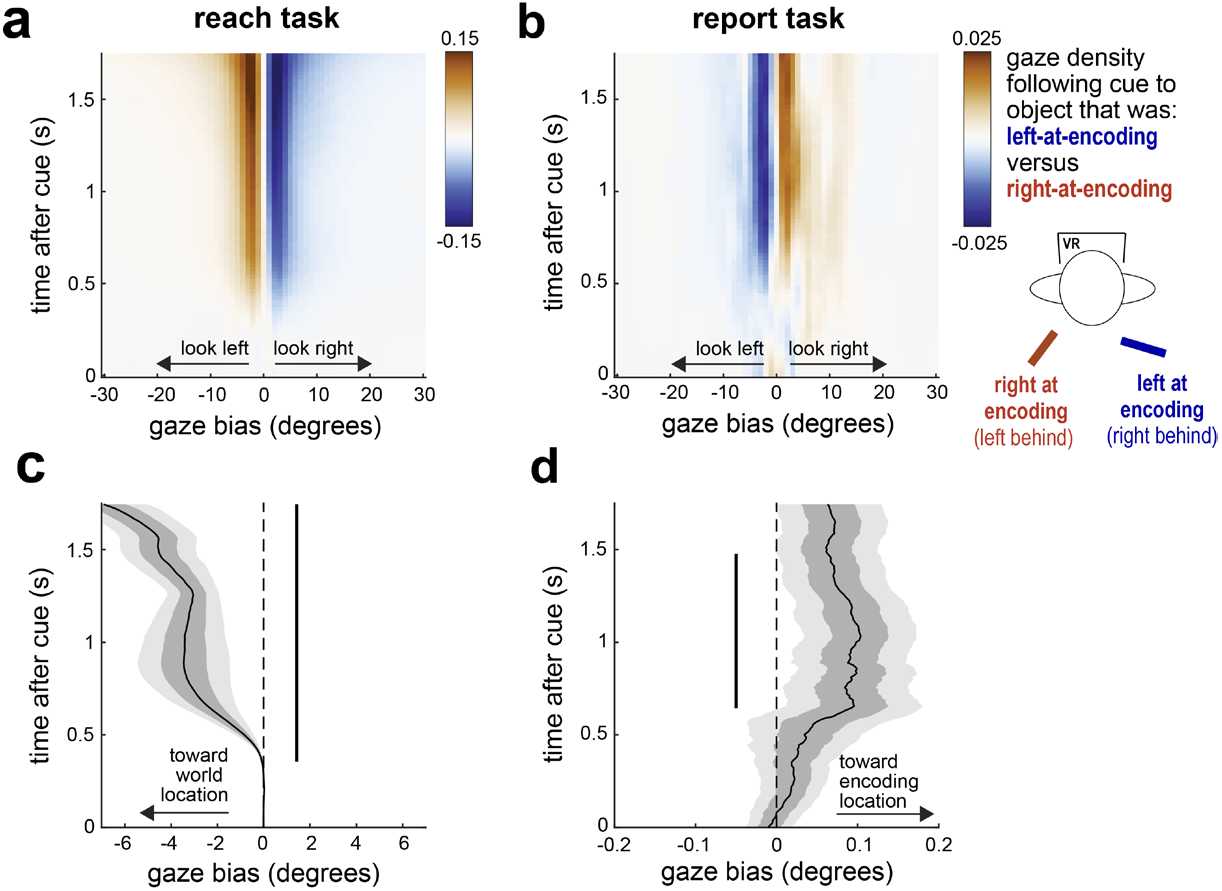
Gaze biases corroborate distinct spatial frames during mnemonic selection when memorising for reaching versus reporting. **a**,**b)** Time-resolved gaze density maps showing the spatial bias associated with cueing the memory object that was left (here indicated in red) vs. right (here indicated in blue) at encoding in the reach task (panel a) and the report task (panel b). The colourmap is chosen to match the schematic of the example trial on the right (see text for further elaboration). **c**,**d)** Gaze-bias towardness time courses, with negative values representing gaze bias toward the cued object’s location in the world behind the participant (body-centred) and positive values representing gaze gaze toward the cued object’s location as registered during encoding (eye-centred). Vertical black bars indicate significant clusters (cluster-based permutation test; ^46^).

When considering object selection in the reach task (**Fig. 3a**), participants’ gaze became biased in the direction of the location of the object as considered in the world behind them – after all, the participants’ task was to reach to this object when receiving the upcoming go cue. Specifically, when selecting the red object (in the example trial configuration depicted in **Fig. 3**) that was now to the left behind them, gaze would become biased to the left, even if the object’s native location during encoding (in eye-centred coordinates) was to the right (before rotation). This was confirmed statistically using cluster-based permutation analysis as a temporal cluster with a bias toward the object’s memorised location as defined in the updated spatial frame representing the location of the object behind the participant (**Fig. 3c**; cluster P < 0.001). This is in line with the above described observation that participants eventually turn back in the correct direction, and shows how this “updated location” is already present and engaged early after the memory cue during initial mnemonic selection.

The crucial insight again comes from the report trials, where we found the *opposite* gaze bias (**Fig. 3b**). This time, when prompted to select the red object (in the example trial configuration depicted in **Fig. 3**) that was the same right object at encoding, gaze would not be biased to the left location of the object in the world behind the participant (as in reach trials), but to the right. This is consistent with the behavioural performance data (here shown for object location instead of orientation), and signals the use of a snapshot-like (eye-centred) representation that participants carried forward with them during rotation. This was again confirmed with a statistical cluster, with a bias toward the opposite direction (**Fig. 3d**; cluster P = 0.008).

Strikingly, we found this spatial gaze bias despite never asking participants about the memorised location of the object in report trials, and despite the fact that there was nothing to the left or right to look at after the central colour cue that prompted selection of the appropriate memoranda. Accordingly, this bias in report trials indexes the incidental spatial organisation in visual working memory that is used by the brain even when location is never asked about (i.e., using space as a scaffold for memory organisation, to putatively support object individuation and selection).

These gaze data this provide critical converging evidence for the use of distinct spatial frames as a function of the anticipated memory task, and show how this task-dependent memory organisation is already in place at the time of mnemonic selection.

## Discussion

By studying visual working memory in moving humans in a controlled VR environment, our data reveal how anticipated action goals fundamentally sculpt the way in which we organise and retain information in visual working memory following self movement – using distinct spatial organisations when anticipating to reproduce vs. reach back to memorised visual objects. We showed this while using identical visual information across tasks. Thus, anticipated memory demands not only shape what aspects of visual information we remember (e.g., ^6,9–14,21,47^), but can also steer us to remember the same visual features (here, location and orientation) in a different way: using a different spatial organisational format. By asking participants to turn 180 degrees, we could uniquely oppose and disentangle two distinct formats to organise visual information in working memory and interrogate the employed spatial organisation through orientation reports and directional gaze biases associated with mnemonic selection. These two sources of evidence converged on our key conclusion that our anticipated behavioural goals fundamentally structure how we hold information in visual working memory following self movement.

In our reach task, we found that participants updated the spatial organisation in memory to represent where the objects were in the world behind them, after having turned. This finding is intuitive, provided that their task was to reach back to the object at its world location. The main insight comes from the report task where we observed evidence for the use of distinct (in our study opposite) spatial memory organisation. In this task, orientation reports consistently adhered to a snapshot-like (eye-centred) representation from the orientation as observed at encoding. Consistently, our spatial gaze marker of mnemonic selection occurred in the opposite direction, again consistent with a snapshot-like representation (i.e. keeping the left-encoded object “left” in memory, despite it physically being to the right in world behind the participant after rotation). Of course, an important difference between our reach and report tasks is that the reach task required keeping track of object location while the report task did not. However, note that in the report task – in which we *never* asked about object location – participants did not simply forgot the memorised object location. Indeed, we also observed a spatial gaze bias in this task, just in the opposite direction. Thus, our key findings do not regard whether our tasks recruit spatial organisation or not, but rather demonstrate how separate spatial organisations can be flexibly recruited depending on the anticipated memory-task.

An interesting question pertains to why participants may have naturally chosen to rely on the snapshot-like (eye-centred) representation in the continuous-reproduction task (a popular working-memory task in the literature; e.g. ^17,19,48,49^). We speculate that participants may have relied on this representation by default, simply because retaining information in this native format required no updating according to self movement. Moreover, there is evidence that memories retained in this native retinal frame may be more precise ^40^. In relation to this specific aspect of our data, it is also interesting that from the 50 participants we tested, only 6 realised that the report could potentially be performed in multiple frames and asked the experimenter what to do. In these cases, we instructed participant to respond in the way that felt more natural to them.

Our findings extend prior research on spatial reference frames in perception and action where it has been well established how vision and action engage multiple reference frames ^28–36^ that, moreover, can be deployed in a task-dependent manner (e.g., ^37,50–52^). Our findings stand out by uniquely addressing such goal-dependence use of distinct frames for memory organisation when retaining and selecting visual information in working memory (and without ever explicitly probing the employed spatial frame). While several prior studies also studied reference frames in working memory ^32,39,40,42^, in this literature it has been common to ask participants where something was, thus engaging explicit spatial memory. A key feature of our critical report task was that we were able to deduce the spatial frame of reference without ever explicitly asking where something was remembered. This was the case both for the orientation-report data (where the “correct” response is mirror-symmetric between the two potential frames), as well as for the spatial biases in gaze behaviour that we measured as a peripheral readout of internal focusing in memory, and of which participants likely remained unaware.

To describe the spatial format (frame) in which participants update and remember the object’s location and orientation as it exists in the world behind them, we referred to “updated” and “body-centred”. We used these terms to refer to a frame that is distinct from the native retinal frame in which the objects were registered at encoding (prior to rotation). While representations in the encoding-centred frame require no additional spatial transformation when moving, keeping track of where the objects are in the world requires these representations to be updated according to one’s own movements through the world. In our study, we got participants to turn 180 degrees which uniquely yielded mirror symmetric orientation and locations in the two frames that we leveraged here. Further research is required to break down the the precise nature of our updated (body-centred) frame (e.g., to carefully delineate whether this is primarily anchored to the current facing direction of the head or the body). Even if our data do not allow to resolve the precise anchoring of this updated frame, our data provide clear evidence for *distinct* spatial frames used for organising memory following movement, driven by the anticipated action goals.

The use of eye movements to interrogate the spatial organisation used for visual working memory builds on a series of recent studies employing a similar approach ^17,25,43–45^ (see also ^23,53–55^). For example, using eye-movements we have recently shown how participants may retain visual information with regard to both its last-seen (past) location and its anticipated (future) location ^6^; how space remains an important organising principle even when temporal order is required for mnemonic object individuation ^25^; or how participants retain spatial organisation in memory even for objects that disappear behind them through optic flow ^44^. Furthermore, in a recent study that we directly built on here, we studied spatial frames used for visual working memory following 90-degree body rotation ^38^. There, we reported evidence for joint reliance on the two spatial frames that we pit against each other here. By using 180-degree rotation in the current study, we uniquely put the frames in competition and additionally manipulated the working-memory tasks to see whether distinct frames would predominate in the distinct tasks. We wish to point out, however, that we do not think the use of one frame or another is necessarily all-or-none. In situations where we rotate more or less than 180 degrees, the two studied frames do not compete, and we may rely on multiple frames concurrently, compatible with our prior findings following 90-degree rotation ^38^.

By studying visual working memory in moving participants, we open new questions regarding the organising principles used for memory when we move through the world. Moreover, by rotating participants 180 degrees, we found a powerful way to infer the employed spatial frame through performance on an orientation-reproduction task (by virtue of the mirror symmetry that emerges between the two frames after rotating 180 degrees), as well as through the direction of spatial biases in gaze during mnemonic selection. These approaches can be taken forward in future research by additionally manipulating other relevant memory features such as set-size, the nature of the stimuli, and object depth; and by testing additional tasks and complementary body movements. Moreover, it may prove interesting to adopt our approach to study cohorts with memory of attentional deficits, and across development and ageing. In addition, it will be interesting to interrogate the neural basis of task-dependent spatial organisation in working memory, such as the contributions of the dorsal and ventral streams ^56–58^. Our experimental approaches and findings open the door to such future opportunities, and provide novel impetus to the wider movement of studying human memory “in action” ^59–65^.

## Methods

### Ethics

The study was conducted in accordance with guidelines approved by the local Research Ethics Committee at the Vrije Universiteit Amsterdam. Participation occurred on voluntary basis, after providing written informed consent. Participants were reimbursed 10 euro/hour or were assigned study credits (each participant selected one of the two types of participation reward).

### Participants

50 healthy adult volunteers participated in the experiment. Of the sample, 18 self-identified as male, 31 self-identified as female, and 1 self-identified as non-binary. The age range was 18 to 29. All participants were right handed. Participants had no known history of neurological or psychiatric conditions, no history of VR-related motion-sickness, and had normal or corrected-to-normal vision. Participants were recruited through an online study registration system (Sona Systems) and through convenience sampling.

Sample size was set a-priori. In prior studies from our lab with similar gaze outcome measures (e.g., ^17,38,44^) we had adopted a sample size of 25. Because we here adopted two task variants per participant (completed in two consecutive sessions, in counterbalanced order), we decided to double the sample size to 50. To reach this intended sample size, 9 participants had to be replaced due to VR-related nausea, or terminating their voluntary participation prematurely, such as after having only completing the first of the two consecutive sessions.

### Design and procedure

Participants performed a visual working-memory task in an immersive VR environment (**Fig. 1**). In different sessions, participants either reproduced the precise orientation of the cued visual object, or reached back to grab it. Importantly, in both tasks, a response was prompted only after having turned their backs to the objects, following a 180-degree rotation. This 180-degree rotation uniquely enabled us to disentangle two potential spatial frames that participants may use for memorising the objects: the native eye-centred frame, containing where the objects are remembered inside a picture-like mental snapshot of the visual objects as registered during encoding prior to rotation, and an updated body-centred frame, keeping track of where the objects are in the external world around (behind) oneself relative to one’s current facing direction. For example, an object encoded to the left before rotation remains the left-side object after rotation in the picture-like mental representation (‘left in the snapshot’); however, this same object could also be remembered as the right-side object after rotation, from the updated frame of reference (‘at the right behind me given my current facing direction’). We investigated the reliance on these two potential spatial frames as a function of the anticipated task: requiring orientation reproduction (report task) or manual grasping (reach task). To this end, every participant completed two experimental sessions, one in which they were always required the report back the orientation of either of two visual objects (that was cued after rotation), and one in which they were always prompted to reach back to the cued memory object and manually grasp it.

Trials in the experiment (**Fig. 1** for a schematic) started with the presentation of the stimuli to the left and right of the central fixation marker. 1000 ms after stimuli onset, the fixation dot would orbit 180 degrees around the participant (taking 3000 ms). Participants were instructed to follow the fixation dot and rotate along with it. 1000 ms after the fixation dot had landed in its new location, the fixation dot would change colour for 1000 ms to match that of either memorised object designating this memory object as the target object for the upcoming task – and thus prompting its selection from working memory. To ensure the use of the cue, the fixation dot would return to its original grey colour for 500 ms before the final memory test.

To simulate a more immersive experience, the bars never disappeared from the environment, but only became ‘out of view’ – and thus to be memorised – because of participants’ own body rotation. However, to ensure the use of the colour cue for selecting the correct memory object, we did remove the bar colour once the bars were no longer in view, by turning both bars black after completing the 180-degree rotation.

The type of memory test was the only thing that differed between our two key experimental sessions. In the reach session, a guiding text (reading “grab”) would appear above the fixation marker, prompting the participant to manually reach back to and grasp the cued memory object with their virtual hand, aiming to grab it in the appropriate orientation. In contrast, in the report session, a circular dial would appear around the fixation marker, whose rotation could be controlled by the hand-held controller. In this case, participants were asked to reproduce the precise memorised orientation of the cued memory object. We never informed participants that there are multiple potential frames in which orientation could be reported, nor instructed them that one frame was preferred over another. For the minority of participants (n=6) who posed this as a question to us, we simply responded they should report however felt more intuitive to them.

In the reach session, feedback was presented when touching the incorrect object or using the incorrect hand. In both sessions, we also provided feedback when the response took longer than the permitted duration (1500 ms in report trials; 2500 ms in reach trials). In theses cases, red text would appeared at the center of the visual field, informing the participant on the specified erroneous behaviour (“wrong hand”, “too slow”). The next trial would always start 1000 ms after feedback offset, starting from the current facing position. As such, participants alternately faced the ‘front’ wall and the ‘back’ wall at the start of every trial.

The experiment contained 10 blocks of 40 trials each, with the first 5 blocks containing exclusively trials of one experimental condition (either ‘report’ or ‘reach’) and the next 5 blocks of the other experimental condition. This condition ordering was counterbalanced across participants. Before each session, participants performed 40 practice trials in order to familiarise with the VR setting and the task.

### VR set-up and stimuli

The experiment was conducted in a physical lab space of approximately 5 x 8 x 4 (h) meter. The VR experience was enabled by an HTC Vive Pro Eye head-mounted display with built-in eye-tracker, an additional Vive Wireless module (allowing increased freedom of movement during VR), two hand-held Vive controllers (for interaction), and two base stations (for spatial tracking). The VR setup was powered by a desktop computer with a NVidia Quadro RTX 4000 graphics processor. The VR environment was developed in Unity (2020.3.9f1) software, and utilised SteamVR (2.6.1) as locational tracking interface and SRanipal (1.1.0) as eye-tracking data interface.

The participant was to stand in the middle of the room, facing the ‘front’ (long) wall, with the two motion-tracking base stations respectively positioned in the two corners of the room in front of them (one base station on the floor, the other mounted on its appropriate stand at a height of 1.5 m), and pointed towards the center of the room. The conducting researcher remained at the desk on one side of the room, with sufficient space around the participant for potential movement or displacement. Where necessary, the conducting researcher could provide additional instructions for the participant in case of difficulties interacting with the VR setup.

The virtual environment was modeled as a plain cubicle of 10 x 10 x 5 (h) meter. The inner texture of the room was homogeneously white (notwithstanding sources of light), with a black line on the floor of the virtual room of 1 m long and 10 cm wide marking the center of the virtual room (providing a reference for the participant in case their standing location would shift over time). A small floating grey sphere at participant eye-height served as the central fixation marker for the participant. This marker also guided the 180-degree rotation. Any instructions that were communicated while the participant was inside the VR environment appeared as text on the (virtual) wall that was expected to be in front of the participant. The participant could interact with the experimental procedure by means of a trigger-like button on the hand-held controllers of the VR equipment.

The critical (controlled) stimuli in every procedural trial of the experiment were two differently-coloured opaque cylinders (with a length of ∼40 cm and diameter of ∼5 cm), each presented respectively 50 cm to the right and to the left of the fixation marker, all at a distance of ∼2 m from the participant. Possible object colours were red, green, blue, cyan, magenta, and yellow. Object orientations (on the plane perpendicular to the participant’s gaze direction) were drawn randomly, with a minimum angular distance of 5 degrees from either the horizontal or vertical axis.

### Analysis

To study the spatial organisation used for visual working memory in our two tasks, we relied on two complementary outcome measures: the orientations associated with orientation-reproduction reports, and spatial biases in gaze during mnemonic selection in both tasks. We discuss each in turn.

For the behavioural-performance data, we leveraged the fact that after rotating around 180 degrees, object orientations can be remembered in two distinct ways. Imagine encoding an object with a leftward tilted orientation (“\”) and rotating around. After rotation, you may remember the object precisely as you have seen it (“\”; i.e. as if carrying a picture of it with you), or update your representation to match the object’s orientation in the world behind you, which is now tilted in the opposite way (“/”) relative to your body. For each orientation-reproduction report, we could thus quantify the error relative to orientation as seen during encoding (native eye-centred frame), or to its mirror image (updated body-centred frame to match the object in the world behind the participant, after 180-degree body rotation).

We visualized response densities relative to both frames in 10-degree bins, that we moved across error ranging from -85 to +85 degree in steps of 1 degree. For statistical quantification, we calculated the absolute error of the report to the orientation of the cued memory object with regard to both frames. We then used a paired-samples t-test to compare frames with regard to participant-specific trial-averaged absolute reproduction errors.

For the gaze data, we similarly leveraged the fact that after rotating around 180 degrees, a left/right object can be memorised in multiple ways. This time consider the object that is to left of the participant at encoding. In the native eye-centred frame – representing a picture-like snapshot of the objects as registered during encoding – the object remains represented as the left object, even after rotation. In contrast, in the updated body-centred frame – keeping track of where the object is in the world behind the participant relative to the participant’s current facing direction – the left object at encoding becomes represented as the object to the *right* behind the participant. To provide complementary evidence for the spatial organisation used for working memory following rotation in our two tasks, we therefore also turned to a gaze signal that we could measure at an earlier stage in the trial: directional biases in gaze signalling mnemonic selection within the spatial-layout of working memory (building on: ^17,25,43–45^). We could do this in both tasks.

To quantify spatial biases in gaze during mnemonic selection, we again first quantified gaze density following cues to objects that were left or right at encoding. Gaze density was calculated in a time-resolved manner, using 200 ms windows that were advanced over the data in 20 ms steps. Density was calculated for the horizontal angle of gaze, in 2-degree bins that were moved over the data from -100 to +100 degrees in steps of 1 degree (with 0 denoting looking straight ahead after rotation). Density values were then subtracted between trials with cues to objects that were left versus right at encoding, to expose any spatial bias associated with mnemonic selection. In addition to these density visualisations, we quantified the gaze ‘towardness’ (as in ^17,25,38,44^) as the average deviation in gaze toward the memorised location of the cue object. We expressed this as a negative value when gaze became biased to the location in the world behind the object, and as a positive value when gaze became biased in the direction of the object as seen during encoding prior to rotation. These towardness time courses were statistically evaluated by means of a cluster-based permutation test ^46^. In this approach, we first clustered neighboring timepoints that reached univariate significance in a one-sample t-test with a alpha-level of 0.05. These clusters were then evaluated under a single permutation distribution of the largest cluster observed after randomly sign-flipping the participant-specific towardness time courses. We evaluated clusters in the observed data to permutation distributions with 10.000 permutations, and calculated p values as the proportion of clusters after permutation that were equal to or larger than the clusters observed in our original data.

Finally, for the reach trials, we quantified how often participants would turn back in the correct direction to grab the cued object behind them (e.g., if the cued object was to the right behind them, how often would participants turn to the right after receiving the go-cue). For this, we calculated the proportion of trials where the average orientation of the VR head-set was deviated more toward the correct (shortest path to cued object) or more towards the opposite (longest path to cued object) direction in the 1000 ms following the go (“grab”) signal.

## Data and code availability

Data and code will be made publicly available prior to publication.

## Notes

### Competing Interest Statement

The authors have declared no competing interest.

### Funding

- European Research Council: 850636
- NWO: 14721

## References

1. Miller, G. A., Galanter, E. & Pribram, K. H. Plans and the Structure of Behavior. (Henry Holt and Company, New York, 1960).

2. Baddeley, A. Working memory. Science (1979) 255, 556–559 (1992).

3. D’Esposito, M. & Postle, B. R. The cognitive neuroscience of working memory. Annu Rev Psychol 66, 115–142 (2015).

4. van Ede, F. & Nobre, A. C. Turning attention inside out: How working memory serves behavior. Annu Rev Psychol 74, 137–165 (2023).

5. Warden, M. R. & Miller, E. K. Task-dependent changes in short-term memory in the prefrontal cortex. Journal of Neuroscience 30, 15801–15810 (2010).

6. Liu, B., Alexopoulou, Z. & van Ede, F. Jointly looking to the past and the future in visual working memory. Elife 12, RP90874 (2023).

7. Christophel, T. B., Iamshchinina, P., Yan, C., Allefeld, C. & Haynes, J. D. Cortical specialization for attended versus unattended working memory. Nat Neurosci 21, 494–496 (2018).

8. Olivers, C. N. L., Peters, J., Houtkamp, R. & Roelfsema, P. R. Different states in visual working memory: When it guides attention and when it does not. Trends Cogn Sci 15, 327–334 (2011).

9. Kwak, Y. & Curtis, C. E. Unveiling the abstract format of mnemonic representations. Neuron 110, 1822–1828.e5 (2022).

10. Nasrawi, R., Boettcher, S. E. P. & van Ede, F. Prospection of Potential Actions during Visual Working Memory Starts Early, Is Flexible, and Predicts Behavior. The Journal of Neuroscience 43, 8515–8524 (2023).

11. Serences, J. T., Ester, E. F., Vogel, E. K. & Awh, E. Stimulus-specific delay activity in human primary visual cortex. Psychol Sci 20, 207–14 (2009).

12. Lee, S. H., Kravitz, D. J. & Baker, C. I. Goal-dependent dissociation of visual and prefrontal cortices during working memory. Nat Neurosci 16, 997–9 (2013).

13. Cao, R. & Deouell, L. Y. Binding in visual working memory is task dependent. J Vis 25, 4 (2025).

14. Niklaus, M., Nobre, A. C. & van Ede, F. Feature-based attentional weighting and spreading in visual working memory. Sci Rep 7, 42384 (2017).

15. Park, Y. E., Sy, J. L., Hong, S. W. & Tong, F. Reprioritization of features of multidimensional objects stored in visual working memory. Psychol Sci 28, 1773–1785 (2017).

16. Schneegans, S. & Bays, P. M. Neural architecture for feature binding in visual working memory. The Journal of Neuroscience 37, 3913–3925 (2017).

17. van Ede, F., Chekroud, S. R. & Nobre, A. C. Human gaze tracks attentional focusing in memorized visual space. Nat Hum Behav 3, 462–470 (2019).

18. Treisman, A. & Zhang, W. Location and binding in visual working memory. Mem Cognit 34, (2006).

19. Pertzov, Y. & Husain, M. The privileged role of location in visual working memory. Atten Percept Psychophys 76, 1914–1924 (2014).

20. Foster, J. J., Bsales, E. M., Jaffe, R. J. & Awh, E. Alpha-band activity reveals spontaneous representations of spatial position in visual working memory. Current Biology 27, 3216–3223 (2017).

21. Griffin, I. C. & Nobre, A. C. Orienting attention to locations in internal representations. J Cogn Neurosci 15, 1176–94 (2003).

22. Kuo, B.-C., Rao, A., Lepsien, J. & Nobre, A. C. Searching for targets within the spatial layout of visual short-term memory. Journal of Neuroscience 29, 8032–8 (2009).

23. Wynn, J. S., Shen, K. & Ryan, J. D. Eye movements actively reinstate spatiotemporal mnemonic content. Vision 3, 21 (2019).

24. Liu, B., Alexopoulou, Z.-S., Kong, S., Zonneveld, A. & van Ede, F. Sparse spatial scaffolding for visual working memory. Preprint at 10.1101/2023.07.05.547765 (2023).

25. de Vries, E., Fejer, G. & van Ede, F. No obligatory trade-off between the use of space and time for working memory. Communications Psychology 1, (2023).

26. Jiang, Y., Olson, I. R. & Chun, M. M. Organization of visual short-term memory. J Exp Psychol Learn Mem Cogn 26, 683–702 (2000).

27. de Vries, E. & van Ede, F. Microsaccades reveal preserved spatial organisation in visual working memory despite decay in location-based rehearsal. Cognition 259, 106111 (2025).

28. Duhamel, J. R., Colby, C. L. & Goldberg, M. E. The updating of the representation of visual space in parietal cortex by intended eye movements. Science (1979) 255, 90–92 (1992).

29. Jiang, Y. V. & Swallow, K. M. Spatial reference frame of incidentally learned attention. Cognition 126, 378–390 (2013).

30. Beurze, S. M., Van Pelt, S. & Medendorp, W. P. Behavioral Reference Frames for Planning Human Reaching Movements. J Neurophysiol 96, 352–362 (2006).

31. Fiehler, K. & Karimpur, H. Spatial coding for action across spatial scales. Nature Reviews Psychology 2, 72–84 (2022).

32. Shelton, A. L. & McNamara, T. P. Systems of Spatial Reference in Human Memory. Cogn Psychol 43, 274–310 (2001).

33. Burgess, N. Spatial memory: how egocentric and allocentric combine. Trends Cogn Sci 10, 551–557 (2006).

34. Golomb, J. D. & Mazer, J. A. Visual Remapping. Annu Rev Vis Sci 7, 257–277 (2021).

35. Tatler, B. W. & Land, M. F. Vision and the representation of the surroundings in spatial memory. Philos Trans R Soc Lond B Biol Sci 366, 596–610 (2011).

36. Pertzov, Y., Avidan, G. & Zohary, E. Multiple Reference Frames for Saccadic Planning in the Human Parietal Cortex. The Journal of Neuroscience 31, 1059–1068 (2011).

37. Sasaki, R., Anzai, A., Angelaki, D. E. & DeAngelis, G. C. Flexible coding of object motion in multiple reference frames by parietal cortex neurons. Nat Neurosci 23, 1004–1015 (2020).

38. Draschkow, D., Nobre, A. C. & van Ede, F. Multiple spatial frames for immersive working memory. Nat Hum Behav 6, 536–544 (2022).

39. Klinghammer, M., Schütz, I., Blohm, G. & Fiehler, K. Allocentric information is used for memory-guided reaching in depth: A virtual reality study. Vision Res 129, 13–24 (2016).

40. Golomb, J. D. & Kanwisher, N. Retinotopic memory is more precise than spatiotopic memory. Proceedings of the National Academy of Sciences 109, 1796–1801 (2012).

41. Postle, B. R. & D’Esposito, M. Spatial working memory activity of the caudate nucleus is sensitive to frame of reference. Cogn Affect Behav Neurosci 3, 133–144 (2003).

42. Aagten-Murphy, D. & Bays, P. M. Independent working memory resources for egocentric and allocentric spatial information. PLoS Comput Biol 5, e1006563 (2019).

43. Liu, B., Nobre, A. C. & van Ede, F. Functional but not obligatory link between microsaccades and neural modulation by covert spatial attention. Nat Commun 13, 3503 (2022).

44. Chawoush, B., Draschkow, D. & van Ede, F. Capacity and selection in immersive visual working memory following naturalistic object disappearance. J Vis 23, 9 (2023).

45. Wang, S. & van Ede, F. Re-focusing visual working memory during expected and unexpected memory tests. Preprint at 10.7554/eLife.100532.1 (2024).

46. Maris, E. & Oostenveld, R. Nonparametric statistical testing of EEG- and MEG-data. J Neurosci Methods 164, 177–90 (2007).

47. Rainer, G., Rao, S. C. & Miller, E. K. Prospective coding for objects in primate prefrontal cortex. The Journal of Neuroscience 19, 5493–5505 (1999).

48. Ma, W. J., Husain, M. & Bays, P. M. Changing concepts of working memory. Nature Neuroscience Preprint at 10.1038/nn.3655 (2014).

49. Zhang, W. & Luck, S. J. Discrete fixed-resolution representations in visual working memory. Nature 453, 233–235 (2008).

50. Pritchett, L. M., Carnevale, M. J. & Harris, L. R. Reference frames for coding touch location depend on the task. Exp Brain Res 222, 437–445 (2012).

51. Ghafouri, M., Archambault, P. S., Adamovich, S. V & Feldman, A. G. Pointing movements may be produced in different frames of reference depending on the task demand. Brain Res 929, 117–128 (2002).

52. Taylor, H. A., Naylor, S. J. & Chechile, N. A. Goal-specific in?uences on the representation of spatial perspective. Mem Cognit 27, 309–319 (1999).

53. Johansson, R. & Johansson, M. Look here, eye movements play a functional role in memory retrieval. Psychol Sci 25, 236–242 (2014).

54. Ferreira, F., Apel, J. & Henderson, J. M. Taking a new look at looking at nothing. Trends Cogn Sci 12, 405–410 (2008).

55. Spivey, M. J. & Geng, J. J. Oculomotor mechanisms activated by imagery and memory: Eye movements to absent objects. Psychol Res 65, 235–241 (2001).

56. James, T. W., Humphrey, G. K., Gati, J. S., Menon, R. S. & Goodale, M. A. Differential Effects of Viewpoint on Object-Driven Activation in Dorsal and Ventral Streams. Neuron 35, 793–801 (2002).

57. Goodale, M. A. & Milner, A. D. Separate visual pathways for perception and action. Trends Neurosci 15, 20–5 (1992).

58. Xu, Y. The Posterior Parietal Cortex in Adaptive Visual Processing. Trends Neurosci 41, 806–822 (2018).

59. Cisek, P. & Green, A. M. Toward a neuroscience of natural behavior. Curr Opin Neurobiol 86, 102859 (2024).

60. Engel, A. K., Maye, A., Kurthen, M. & König, P. Where’s the action? The pragmatic turn in cognitive science. Trends Cogn Sci 17, 202–209 (2013).

61. Olivers, C. N. L. & Roelfsema, P. R. Attention for action in visual working memory. Cortex 131, 179–194 (2020).

62. van Ede, F. Visual working memory and action: Functional links and bi-directional in?uences. Vis cogn 28, 401–413 (2020).

63. Heuer, A., Ohl, S. & Rolfs, M. Memory for action: a functional view of selection in visual working memory. Vis cogn 28, 388–400 (2020).

64. Stangl, M., Maoz, S. L. & Suthana, N. Mobile cognition: imaging the human brain in the ‘real world’. Nat Rev Neurosci 24, 347–362 (2023).

65. Draschkow, D., Kallmayer, M. & Nobre, A. C. When natural behavior engages working memory. Current Biology 31, 869–874.e5 (2021).

